# Population cycles and species diversity in dynamic Kill-the-Winner model of microbial ecosystems

**DOI:** 10.1101/067124

**Authors:** Sergei Maslov, Kim Sneppen

## Abstract

Determinants of species diversity in microbial ecosystems remain poorly understood. Bacteriophages are believed to increase the diversity by the virtue of Kill-the-Winner infection bias preventing the fastest growing organism from taking over the community. Phage-bacterial ecosystems are traditionally described in terms of the static equilibrium state of Lotka-Volterra equations in which bacterial growth is exactly balanced by losses due to phage predation. Here we consider a more dynamic scenario in which phage infections give rise to abrupt and severe collapses of bacterial populations whenever they become sufficiently large. As a consequence, each bacterial population in our model follows cyclic dynamics of exponential growth interrupted by sudden declines. The total population of all species fluctuates around the carrying capacity of the environment, making these cycles cryptic. While a subset of the slowest growing species in our model is always driven towards extinction, in general the overall ecosystem diversity remains high. The number of surviving species is inversely proportional to the variation in their growth rates but increases with the frequency and severity of phage-induced collapses. Thus counter-intuitively we predict that microbial communities exposed to more violent perturbations should have higher diversity.

## I. INTRODUCTION

An important and largely unsolved question in microbial ecology is what determines the diversity of microbial ecosystems. Indeed, unbridled competition between microbes sharing common resources would eventually limit species diversity not to exceed the number of different nutrient types [1]. Predation by bacteriophages introduces the negative frequency-dependent selection [2–5] which offers the possibility for a dramatically larger species diversity [5]. In the classical Kill-the-Winner (KtW) model of Thingstad [5] virulent phages reduce populations of their susceptible hosts to a low steady state level, which is independent of hosts’ growth rate thus allowing multiple species per nutrient type. The number of co-existing bacterial species in the resulting ecosystem is determined exclusively by the parameters of phage predation [5], the topology of the phage-bacterial infection network [7–9], and the carrying capacity of the environment [4, 6, 8, 9].

Microbial population dynamics is routinely a much more dynamic process than assumed in the traditional steady state KtW model and its variants. For example, in the lab experiments [10] *E. coli* population suffered a dramatic collapse by a factor ~ 10^4^ – 10^5^ caused by a T7 phage infection. Collapse-driven dynamics is common in both natural [11] and man-made [12–15] ecosystems in which bacteria are engaged in the continuous arms race with phages [16–20].

To capture this here we propose and explore a *dynamical* interpretation of Kill-the Winner principle, in which bacterial populations are characterized by periods of competitive exponential growth punctuated by rapid and severe collapses. Larger bacterial populations in our model are proportionally more likely to be infected by phages. Furthermore, in larger and thus denser populations such infections once started are likely to eliminate a sizable fraction of susceptible hosts resulting in a severe collapse in the populations of individual bacterial strains. When viewed over a long period of time any given species would repeatedly cycle between low and high population numbers. Such cyclic dynamics of populations of individual species masked by an approximately constant total population saturated at the carrying capacity of the environment is discussed in the ecological literature as “cryptic cycles” [21–23].

### Model

Consider a number of bacterial species/strains sharing the same environment and competing for the same rate-limiting nutrient defining its carrying capacity. Their populations sizes at time *t* are denoted as *P_i_*(*t*), where *i* = 1, 2, …, *N*. Each of these individual species is exposed to rare but severe collapse events in which its population is suddenly and drastically reduced by a constant factor *γ* ≪ 1. We assume that these collapses happen relative rarely so that the total population of all bacterial species has sufficient time to reach the steady state value given by the overall carrying capacity of the environment. Without loss of generality carrying capacity can be set to 1, so that in between collapses one has 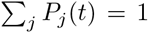. In our model we assume that while the total population of all stays constant, relative population sizes of individual species continue to change exponentially in-between collapse due to differences in their fitness in the saturated environment.

In the spirit of Kill-the-Winner principle we assume that the rate of collapse of the species i is proportional to its population size *P_i_*. Due to a broad distribution of population sizes this rule strongly biases collapses towards one or few largest populations. We assume that collapse events are independent of each other, so that the time interval between consecutive collapses is exponentially distributed with mean *τ*.

One update cycle in our model consist of three steps:

1. Draw a time interval Δ*t* until the next collapse event from the exponential distribution *P*(Δ*t*) = exp(−Δ*t*/*τ*)/*τ*.
2. Calculate population sizes at the time of collapse. In between collapse events relative population sizes are assumed to change exponentially while the total population stays saturated at 1 (the carrying capacity of the environment):

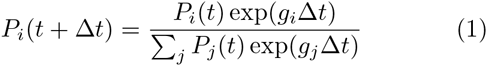
3. Select one species to collapse with the probability equal to its relative population size *P_i_*(*t* + Δ*t*) and multiply its population by *γ*.

In our simulations each of *N* species is assigned its individual growth rate drawn from the Gaussian distribution with zero mean and standard deviation *σ*. The value of the mean is not important since normalization of the overall population to 1 ensures that only relative growth rates matter. Furthermore, the exact form of the distribution of growth rates is not particularly important. In our mathematical analysis we will use a more convenient exponential distribution of growth rates: *P*(*g_i_*) = exp(*g_i_*/*σ*)/*σ*, while delegating more cumbersome derivations for the Gaussian *P*(*g_i_*) to supplementary materials.

## II. RESULTS

### Collapses supports Diversity

Figure 1A shows a typical outcome of a simulation of our model with *γ* = 10^−3^ and *σ* = 4 over around 100 population collapses after which only *D* = 3 out of *N* = 8 species survive. The relative growth rate *g_i_* of the species is the main predictor on whether it will survive or not. Indeed, as shown by the rainbow coloring of curves in Fig. 1A ranging from dark red (the slowest growing) to purple (the fastest growing) the 3 surviving species have the largest values of *g_i_*.

**FIG. 1.**
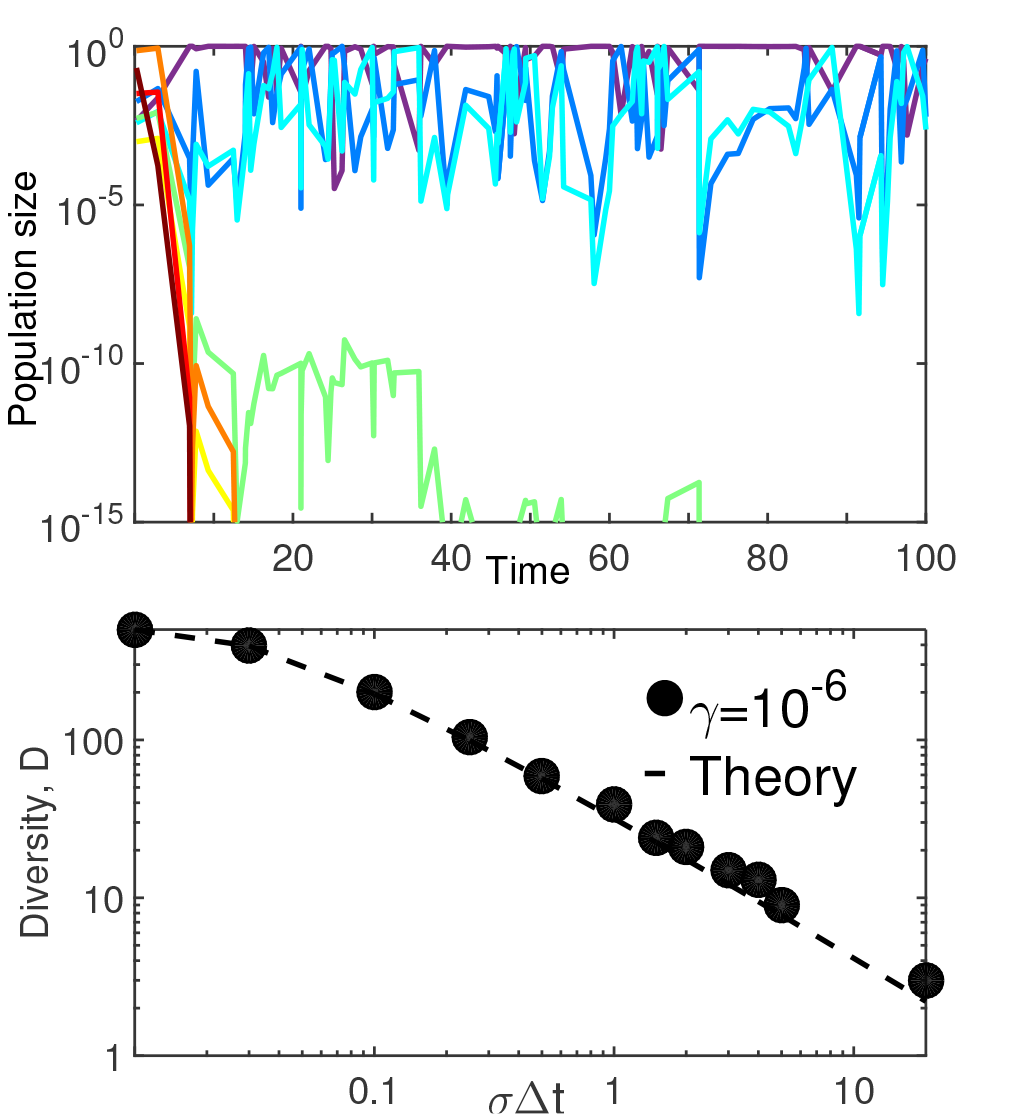
Simulations of Kill-the-Winner model. A) Time courses of populations of N = 8 species with fixed growth rates assigned from the Gaussian distribution with standard deviation σ = 4. Rainbow colors correspond to growth rates ranging from the slowest (dark red) to the fastest (purple). For each of the species, the likelihood of collapse is proportional to its population sizes (“Kill-the-Winner” rule) and the collapse ratio γ = 10^−3^ is the same for all species. Only 3 fastest growing species survive in the long term B) The final diversity D counted as number of surviving species as function of σΔt - the spread of growth rates integrated over the average time between collapses. Each black dot represents the outcome of one simulation started with N = 500 species exposed to collapse ratio γ = 10^−6^. The dashed line is the analytical fit similar to Eq. 4 but here done for the Gaussian distribution of growth rates used in the simulation (see Supplementary Materials for details).

A natural question to ask is what determines the number of surviving species/strains in the steady state of the model?

In the limit of very rare collapses the fastest growing species would diverge from the rest of the population so much that it will be the only one to survive, as indeed expected from the competitive exclusion principle [1].

The situation is more complex for intermediate rate of collapses where more than one of the fastest growing species can coexist with each other but some of the slowest growers become extinct. In the steady state each of these surviving species repeatedly cycles between low and high populations. Faster growing species reach large population sizes more often which makes them to collapse more frequently thus eliminating their growth advantage. As we show below this balance can be sustained within a finite range of growth rates.

For each of the species its individual growth rate *g_i_* is reduced by the same negative number −*g_cc_* due to the overall resource competition quantified by the denominator in Eq. 1. In the steady state the excess growth rate of each of the surviving species (*g_i_* − *g_cc_*) must be exactly compensated by the logarithmic population losses |log *γ*| due to collapses happening at the species-specific probability *c_i_*:

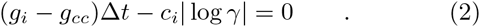

Note that as the probability of collapse (per update) *c_i_* = (*g_i_* − *g_cc_*)Δ*t*/|log *γ*| needs to be positive and normalized. Positivity of *c_i_* means that only the fastest growing species with *g_i_* > *g_cc_* would survive in the long run. The collapse rates of these *D* surviving species are further constrained by normalization 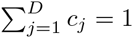, reflecting the requirement of one collapse per update. Using eq. 2 the threshold *g_cc_* is then determined by:

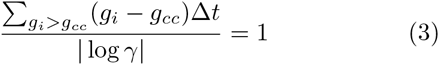

For a given set of species, this allows us to self-consistently calculate *g_cc_* and *D*

For *g_i_* selected from the exponential distribution with standard deviation *σ* the diversity *D* is given by (see Supplementary Information for derivation)

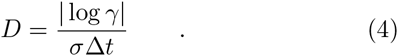

This expression holds for average diversity provided that it is larger than 1 as single a single fastest growing species would always survive. Clearly *D* is also capped from above by *N*. Similar relation holds for the Gaussian distribution of growth rates and is in agreement with our numerical simulations of the model shown in Fig. 1b.

For the exponential distribution the growth rate threshold above which a species survives is given by *g_c_c* = *σ*log(*N*/*D*) = *σ*log(*NσΔt*/|log *γ*|). Note that while threshold explicitly depends on the starting number of species, the final diversity given by Eq. 4 is independent of *N*. This particular property of the exponential distribution would be modified for other distributions resulting in a mild dependence of *D* on *N*.

A Gaussian distribution of growth rates would slightly increase the diversity compared to the exponential distribution with the same spread, while a more fat-tailed *(say, power law) distribution would decrease it.

Our basic model can be generalized to the case where different species have different collapse ratios *γ_i_*. This may for example reflect their different degrees of vulnerability to phages, or different ways to partition their population in physical space. The only consequence of this modification is that log *γ* in the equations above needs to be replaced by its average value across species (see supplementary materials for simulation results).

In our model the collapse probability of a given species is proportional to its population size. Thus time-averaged relative population size of each of the species species is equal to its overall collapse frequency 〈*P_i_*(*t*)〉_*t*_ = *c_i_*. This is consistent with “Kill-the-Winner” principles according to which species with larger populations collapse more often.

Fig. 2b illustrates this cyclic dynamics in a system containing a mixture of slow and fast growing species. Surviving populations mostly grow, but do so at different rates. Their coexistence is possible only because of the negative feedback via “Kill-the-Winner” rule where populations of an individual species get severely reduced once it starts to dominate the overall biomass. The population of each of the species goes through approximately periodic cycles of growth and collapses with the period *T_i_* = 1/*c_i_* = |log *γ_i_*|/(*g_i_* − *g_cc_*)Δ*t* (in units of collapse events). Thus the slowest surviving species (marked blue in Fig. 2b) nearly never collapse, whereas the fastest growing species (marked red in Fig. 2b) obtain dominance and expose themselves to a collapse on a much shorter timescale. Individual collapse events of these species are marked in Fig. 2b with red and blue arrows correspondingly. Note that the population of the slowest growing species often decreases not due to a phage-mediated collapse but simply because it gets temporarily outgrown by other species with a faster growth rate.

**FIG. 2.**
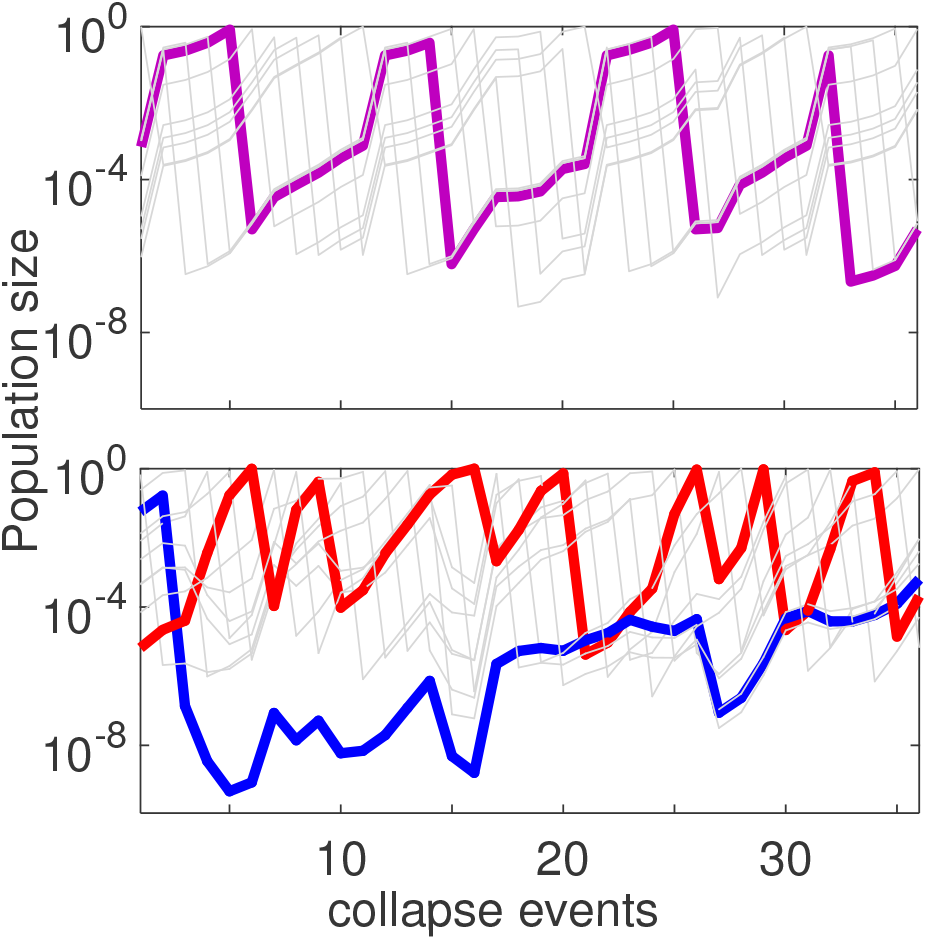
Cyclic dynamics in Kill the Winner model. A) D = 10 species with identical growth rates and, collapse ratios γ_i_ = 10^−6^. The highlighted purple curve illustrates the characteristic growth and collapse cycle for a particular population. Purple arrows indicate collapse events. B) Simulation with D = 10 surviving species (down from N = 60) each with growth rates selected from the Gaussian distribution with σ = 3, and identical collapse ratios γ_i_ = 10^−6^. The blue arrow and the red arrows mark times for collapse events of these two species. Note how the fastest growing species (red line) collapses much more often than the slowest growing species (blue line) which only collapsed once during the time shown. The growth rate difference between these two species is g_max_ – g_min_ =2.44.

For comparison in Fig 2a we show a system of the same size (*D* = 10) but where all species have exactly the same growth rate. In that case the system has a very long memory of the initially imposed order of species populations, because even after a long time each of the species would collapse the same number of times. That is, if one species have experienced one more collapse than the others, it would be smaller by a factor γ and thus be much less exposed to subsequent collapses until it would regrow to the size where it again may collapse with a non-negligible probability. Indeed, populations shown in Fig. 2a follow much more regular oscillatory dynamics than those with unequal growth rates shown in Fig. 2b.

### Model with collapses to a fixed threshold

In our standard version of the KtW model the collapsing population is reduced by a constant factor: *P_i_* → *P_i_* · *γ*. An alternative possibility is that following a collapse the population starts at a fixed small threshold value *γ* irrespective of its earlier population size. This would be the case e.g. when following a collapse the local population is completely eliminated and is reintroduced by one individual from a neighboring region. It can also happen when a collapse drives one species extinct only to be quickly replaced by a single bacterium of a new species. In thus defined fixed threshold kill-the winner model (KtWT) the diversity remains close to what was reported in Fig. 1b (data now shown). The dynamics is also characterized by individual population undergoing cycles of duration defined by their relative growth rates much similar to what is shown in Fig. 2 for our original model. However the long term memory of cycle order is reduced compared to the constant factor model discussed above, simply because every collapse completely erases the population history of the collapsed species. In what follows we explore the dynamical properties of the fixed threshold model and its variants.

### “Kill-the-King” Model

To better understand the cyclic dynamics in the KtWT model we first consider its extreme and deterministic version in which the next collapse always happens at the largest population. We will refer to this version as Kill-the-King (KtK) model. Furthermore, we assume that the growth rates *g_i_* of all species are equal to each other. Thereby the asymptotic dynamics becomes periodic with period *N* when time is measured by collapse events.

To concentrate on slow trends in population size dynamics we only measure them between intervals where each population collapsed exactly once, which in KtK secure that the order of populations is exactly preserved. We relabel species in the order of decreasing population sizes and calculate the ratios *δ_i_* = *P*_*i*+1_/*P_i_* < 1 between successive population sizes [1]. As shown in the supplementary materials, in KtK model these ratios evolve according to the following discrete equation describing changes acquired after a full round of *N* collapses so that each member of the population collapsed exactly once:

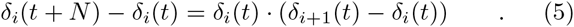

The steady state of the equation is reached when all ratios are equal to each other, i.e. *δ*_*i*+1_ − *δi* = 0. In this case the logarithms of population sizes are equidistantly distributed in the interval of length |log *γ*| so that *δ_i_*(∞) = *δ*\* = *γ*^1/*N*^. Figure 3 shows a simulation of KtK model with *N* =10 and *γ* = 10^−6^ One can see how it asymptotically approaches this steady state.

**FIG. 3.**
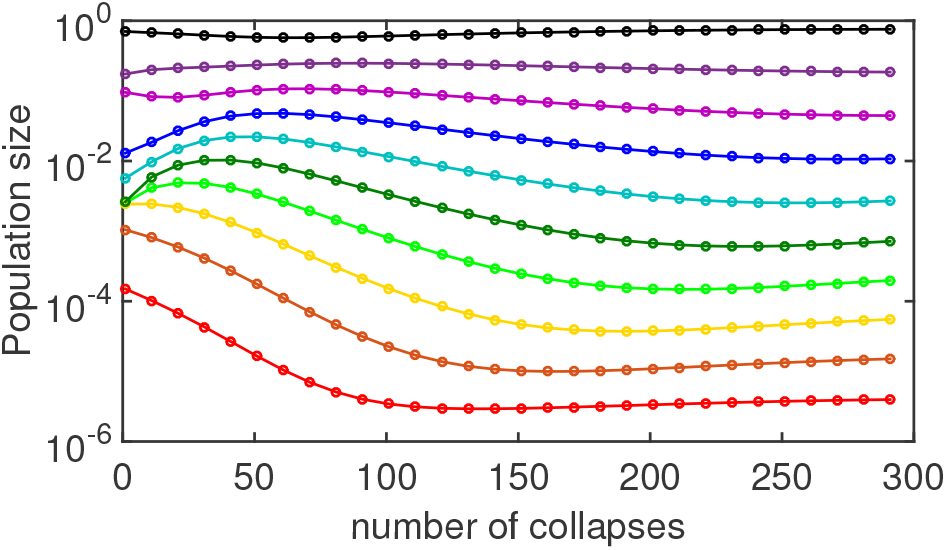
Transient dynamics in Kill-the-King model with N = 10 species that grow equally fast and collapse to a fixed population γ = 10^−6^. For clarity we show the state of the model only every 10 collapses, that is to say, after each species collapsed exactly once so that the population order is maintained. The steady state of KtK model where all ratios δ_i_ between rank-ordered populations are equal to each other and to γ^1/N^ is approximately reached already after 300 collapses. The relaxation to this steady state is described by the discrete anisotropic Burgers equation 5 or its continuous counterpart Eq. 6.

The asymptotic dynamics of KtK is described by the discrete Eq. 5 which for large *N* can be approximated by a continuous PDE (see SI for more details) in which the continuous coordinate *x* replaces the species rank *i*:

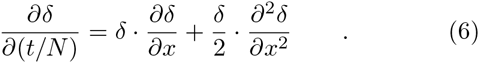

Here *δ*(*x*) has periodic boundary conditions over *x*-interval [0, *N*]. As its discrete counterpart this equation describes the state of our system every *N*’th time-step. This equation is closely related to the Burgers equation [24, 25], although it differs in terms of the diffusion coefficient that instead of being constant as in Refs. [24, 25] is proportional to *δ*(*t*).

Having finished with Kill-the-King model we return to Kill-the-Winner fixed Threshold (KtWT) model. In the KtWT model population collapses do not always happen in the order dictated by their relative sizes. This results in a somewhat chaotic dynamics illustrated in Fig. 4A. When a smaller population collapses out of turn it causes only a very small rescaling of other population sizes. The (very likely) subsequent collapse of the largest population leads to a situation where these two just collapsed populations become nearly equal in size (*δ* ≃ 1). This dramatically increases the likelihood for further re-orderings between these two species, resulting in an extended period where these two species fight for dominance. This intermittent dynamics switching the order of populations is clearly visible in Fig. 4B with *D* = 3. The nearly vertical lines clustered around collapse events 5100, 5200, and 5400 correspond to frequent shifts in the population order of three species within the cycle.

**FIG. 4.**
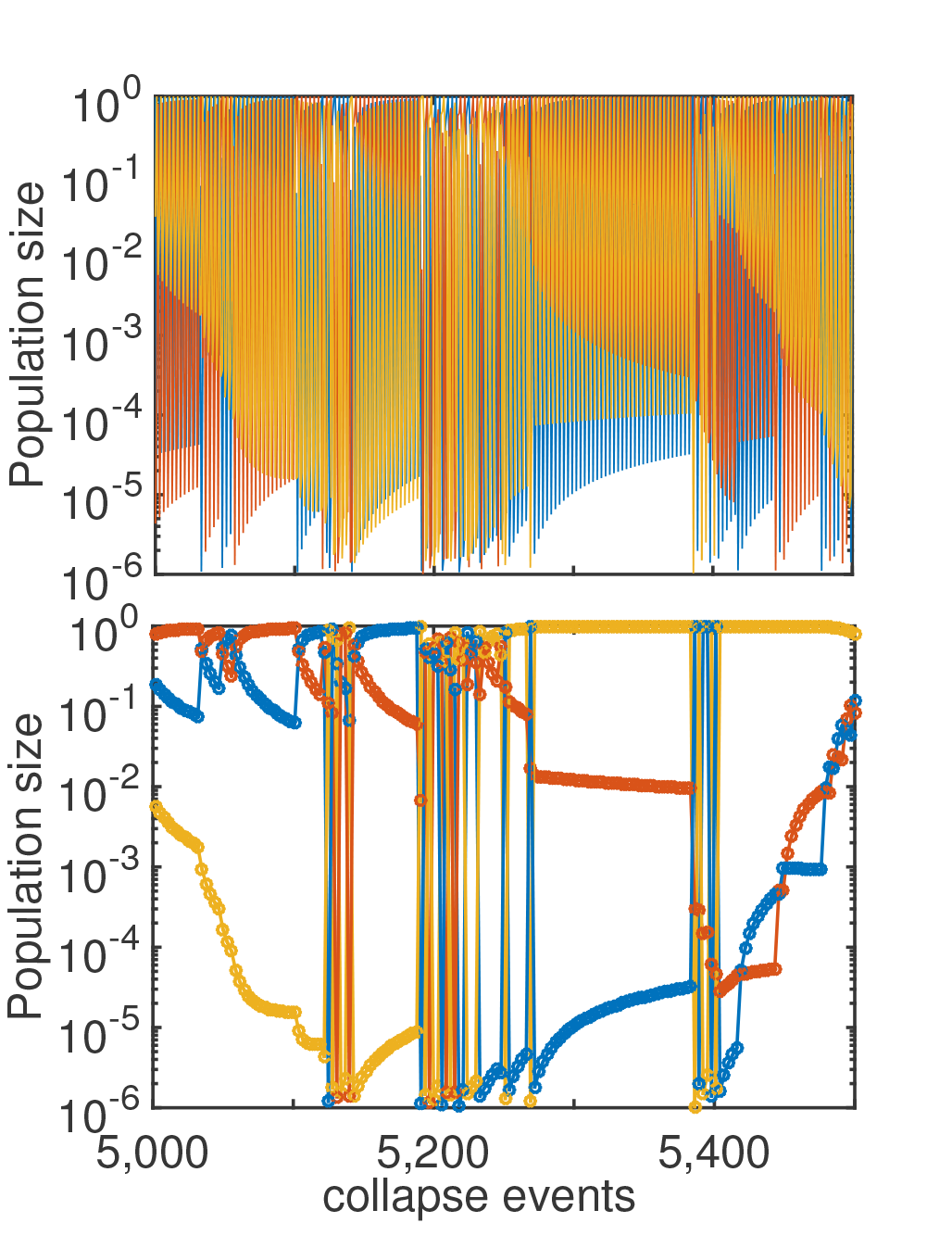
“Kill-the-Winner” Threshold (KtWT) model with 3 species that grow at equal rates and ultimately collapse to a fixed population size γ = 10^−6^. A) Dynamics of all 3 species, emphasizing that the cyclic order occasionally changes, caused by an “out of order” collapse of a population that is not the largest. B) Same time series as in the above panel, but only showing every third time-point. This panel highlights the interplay between occasional intermittent alternations in the cyclic order (clustered vertical lines) and longer “quiet” periods during which, ratios of rank-ordered populations relax towards δ* = γ^1/N^.

An intermittent region ultimately ends with a particular order winning over. After this all populations slowly relax back to the steady state with equal ratios *δ*\* (curved lines ending in horizontal plateaus in Fig. 4B). The exact form of the relaxation to the steady state is derived in supplementary materials. While *δ*(*t*) ≫ *δ*\* the relaxation is proportional to 1/(*t*/*N*). The expected number of collapse events for i*δ*(*t*) to go from ~ 1 to ~ *δ*\* is ~ *N*/*δ*\* or about 300 for the parameters used in Fig. 4. When *δ*(*t*) ≃ *δ*\* the relaxation crosses over to *δ*(*t*) − *δ*\* ~ exp(−*δ*\**t*).

## III. DISCUSSION

Here and before [26] we investigated the impact of severe and sudden population collapses on ecosystem composition and diversity. This approach is complementary to a more traditional description of ecosystem dynamics at or around the steady state solution [2, 5, 6, 28]. The emergent cyclic dynamic in our model is entirely collapse-driven and thus distinct from either stable or transient periodic oscillations present in predator-prey ecosystems described by the Lotka-Volterra equations [22, 23, 27–29].

The key assumption used in our study is that larger populations are more exposed to sudden collapses than the smaller ones This is the foundation of “Kill-the-Winner” (KtW) principle proposed in Ref. [5]). The resulting negative (or stabilizing) frequency-dependent selection promotes the ecosystem diversity even in the simplest case considered above, where species interact with each other only via competition for a single rate-limiting resource. This KtW bias is very important as it shifts the collapse-driven dynamics away from “diversity waves” we reported before [26] towards population cycles investigated in this study. Indeed, as demonstrated in [26] a version of collapse-driven dynamics in which the likelihood of a collapse is uncorrelated with population size or even biased towards smaller populations (Kill-the-Looser model) results in ebbing and flowing species diversity and bi-modal distribution of species abundances. This should be contrasted with time-independent diversity in KtW, KtWT, and KtK models studied here and predictable (at least in the short-term) cycles in which population collapses follow each other in a particular order. The species abundance distribution in these models is not bi-modal but uniform on the logarithmic scale (data not shown).

To test how sensitive are our results with respect to introduction of other types of interactions between bacterial species as well as to a more branched topology of Phage-Bacterial Infection Networks (PBIN) we simulated a variant of our model where in addition to abundant (KtW) species the infecting phage results in collapse of a constant number *K* of other bacterial species. This version of the model is reminiscent of the Bak-Sneppen model of species co-evolution [30]. We tested this model for *K* =1 and *K* = 25 (out of *N* = 500). In the first case we observed no impact on diversity, while in the second case the diversity saturated at lower values of *σ*Δ*t*. All together we can conclude that the diversity profile shown in Figure 1B remains qualitatively (and sometimes even quantitatively) unaffected by additional interactions between microbial species or more interconnected PBINs.

According to our results the principal determinant of the ecosystem diversity *D* is the width *σ* of the distribution of logarithmic growth rates of individual bacterial strains or species. This difference is amplified during the average time Δ*t* between population collapses. Thus the overall frequency of collapses is a very important parameter with more frequent collapses counter-intuitively resulting in more diverse ecosystems. That is because in our scenario frequent collapses weaken the effect of competitive exclusion ultimately driving the diversity down to no more than single species per rate-limiting nutrient. Larger magnitude of collapses also promotes higher diversity but its impact increases only weakly (logarithmically) with the collapse ratio *γ*.

It is instructive to compare the determinants of microbial diversity in the static, steady state KtW model and in our more dynamic, collapse-driven variant. In the static KtW model [2, 5] the steady state population size of each of the bacteria *B*\* = *δ*/*βη* is determined exclusively by parameters of the phage to which it is susceptible: its burst size (*β*), death (or dispersal and dilution) rate (*δ*), and its infection rate (*η*) at a density equal the bacterial carrying capacity. This steady state population of a phage-controlled bacterium is usually much lower than the carrying capacity of the environment: *B*\* << 1. Thus a large number of bacteria each susceptible to its unique phage predator can coexists with each other [5]. Higher diversity can subsequently be achieved by carefully adding pairs of bacteria and phages, latter possibly supplemented by their epigenetic variants [31], each consuming a small fraction of the carrying capacity [5, 6]. Substantial diversity is found to be fragile to new invaders, in the form of bacteria that grow faster than resident ones or phages that prey on several bacteria at once [6].

In contrast to this the diversity in our model is determined by both statistics of collapses as well as the spread of growth rates of resident bacterial species. In case of mild or infrequent collapses and large disparity in bacterial growth rates competitive exclusion principle is restored within our model as it then predict an ecosystem dominated by just one fastest growing bacterium. When collapses are frequent (short Δ*t*) and severe (large |log *γ*|), while growth rates of individual bacterial strains or species are close to each other (small *σ*), Eq. 4 predicts high diversity of co-existing bacterial species. This prediction is robust with respect to exact causes of collapses, including the relatively frequent [15] invasion of phages that are capable of infecting several bacterial species.

Overall, the falsifiable (and counter-intuitive) prediction of the collapse-driven “Kill-the-Winner” model differentiating it from its stationary counterpart, is that by increasing frequency (and to a smaller extent severity) of collapses one could support higher diversity of microorganisms.

## SUPPLEMENTARY MATERIALS

### Diversity for an arbitrary fitness distribution

Let’s consider the case where growth rates *g_i_* is selected from probability distribution with standard deviation *σ*: *P*(*g*) = (1/*σ*)*f*(*g*/*σ*). Here *f*(*x*) is the PDF of the distribution with standard deviation 1. For a large initial population *N* the sum 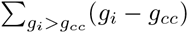 in eq. 3 can be approximated with the integral:

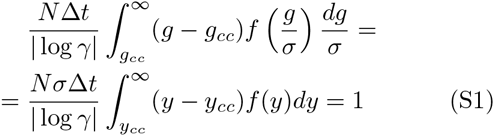

where *y_cc_* = *g_cc_*/*σ* is normalized minimal growth rate needed for a long time survival of the species. The diversity of surviving species is then given by

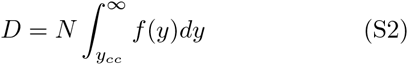

For exponential distribution, *f*(*y*) = *exp*(−*y*), one gets

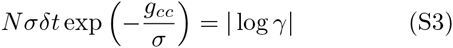

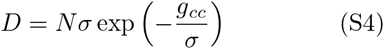

or

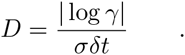

In the case of the Gaussian distribution, 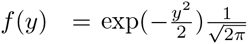, Eqs. ?? cannot be solved analytically. Numerical solution shown as dashed line in Fig. 1b closely resembles simulations.

### Diversity with variable collapse ratios

The model can be directly generalized to the case where different species have different collapse ratios *γ_i_*. This may for example be the case for bacteria with different degrees of vulnerability to phages, or bacteria with different ways to partition their population in physical space. The modified Eqs. 2 and 3 reads

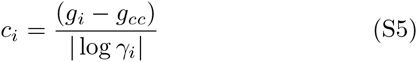

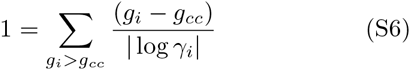

where the first equation again imply that only the *g_i_* > *g_cc_* will contribute.

To solve eq. S6 the sum in eq. S6 is successively tested for species that is rank ordered from the largest values of *g_i_* = *g*_1_, until a value *D* = *i* where it provide a solution for a *g_cc_* ∈ [*g*_*D*+1_, *g_D_*].

Allowing individual species to have different *γ_i_* only moderately changes the diversity *D* compared to the uniform case of *γ_i_* = *γ* (see Fig. S1). We tested several variants in which *γ_i_* was assigned to individual species as described by Eq. S6 or in which *γ_i_* was randomly varying between collapse events. Allowing *γ_i_* to vary between species or collapse events also did not affect the distributions of populations sizes (data not shown).

**FIG. S1.**
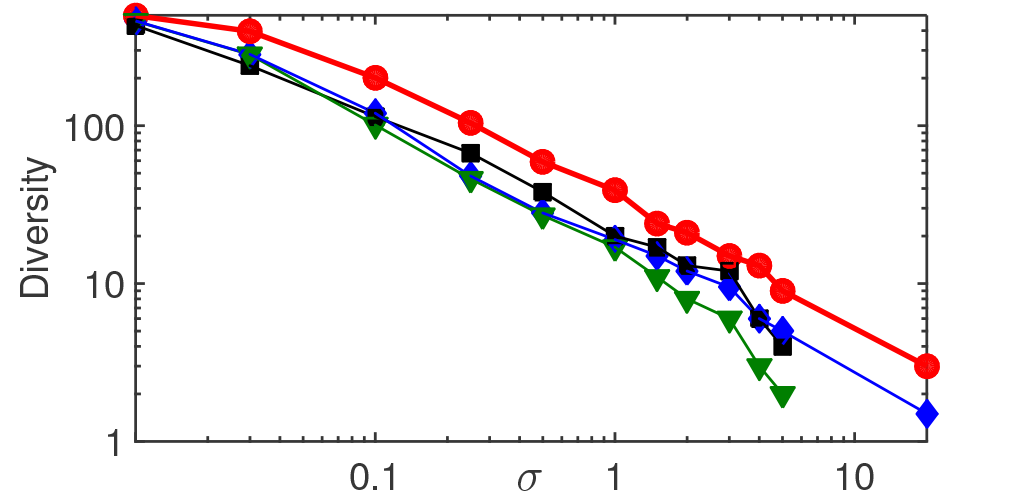
Final diversity D counted as number of species with populations > 10^−20^ as the function of spread σ of initially assigned growth rates among N = 500 species. The red curve marked, with circles corresponds to the KtW model with fixed γ = 10^−6^, while the blue curve marked with diamonds - to fixed γ = 10^−3^. Te black curve marked with squares show simulation results when γ_i_ varies between species (quenched noise), while the red curve marked with triangles - when it varies between collapse events (annealed noise). In these two cases log_10_ γ_i_ was drawn from a uniform distribution between −1 and −5 so that its geometric average of 10^−3^ corresponds to the blue curve.

### Kill-the-King model

For convenience we choose an arbitrary time point *t* = 0 and reorder the populations in the order of decreasing sizes so that *P*_1_ (0) corresponds to the largest population, *P*_2_(0) - to the second largest, and so on with *P_N_*(0) being the smallest population. The new cycle starts with *P*_1_ (0) which collapses down to *γ* while the rest of the species remain unchanged with the total population of 1 − *P*_1_(0). After the collapse all species grow “instantly” to the carrying capacity resulting in *P*_1_(1)= *γ*/(1 − *P*_1_(0) + *γ*).

At the subsequent phage attack the second population *P*_2_(1) collapses which after rescaling leaves the first two populations as respectively

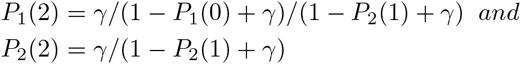

The subsequent *N* − 2 collapses of populations 3 to *N* do not change the ratio between the first two populations, implying that the updated ratio between the second and the first population remains *P*_2_(*N*)/*P*_1_(*N*) = *P*_2_(2)/*P*_1_(2) = 1 − *P*_1_(0) + *γ*.

As derived above then *δ*_1_(*t*+*N*) = 1 − *P*_1_(*t*)+*γ*. Taking into account that 1 = ∑_*i*_ *P_i_* = *P*_1_ + *P*_1_*δ*_1_ + *P*_1_*δ*_1_*δ*_2_ + … *P*_1_*δ*_1_*δ*_2_ … *δ*_*N*−1_, in the limit where all *δ_i_* ≪ 1 up to the second order in *δ_i_* one gets *P*_1_ ≃ 1/(1 + *δ*_1_ + *δ*_1_*δ*_2_) or 1 − *P*_1_ ≃ (*δ*_1_ + *δ*_1_*δ*_2_)/(1 + *δ*_1_ + *δ*_1_*δ*_2_) ≃ (*δ*_1_ + *δ*_1_*δ*_2_)·(1 − *δ*_1_) ≃ *δ*_1_ + *δ*_1_(*δ*_2_ − *δ*_1_). Thus the following equation describes change of *δ* after one full round of population collapses: *δ*_1_(*t*+*N*) = 1 − *P*_1_(*t*)+*γ* ≃ *δ*_1_(*t*)+*δ*_1_(*t*)(*δ*_2_(*t*)−*δ*_1_(*t*)) up to *O*(*δ*^2^). Since in the course of one cycle of collapses each population in turn becomes the largest one, the above equation for *δ*_1_ applies to *i* other than 1:

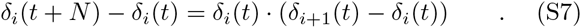

In the long time limit the populations in the KtK model asymptotically approach an equidistant distribution on the logarithmic scale. Thus *δ_i_*(∞) = *δ*\* = *γ*^1/*N*^. Uniform distribution of population sizes on the logarithmic scale corresponds to the power law species abundance distribution *P*(*S*) ~ *S*^−1^.

In the continuous limit in time *t* and space *x* = *i* the Eq. S7 can be rewritten taking into account *δ_i_*(*t* + *N*) − *δ_i_*(*t*) ≃ *N∂δ*(*x, t*)/*∂t*, while *δ*_*i*+1_(*t*) − *δ_i_*(*t*) ≃ *∂δ*(*x, t*)/*∂x*+(1/2)*∂*^2^*δ*(*x, t*)/(*∂x*)^2^. Note the appearance of the second derivative over *x* due to the fact that *δ*_*i*+1_(*t*) − *δ_i_*(*t*) is centered half-way between *i* and *i* + 1 and thus is shifted up from *i* by 1/2. Thus the gap dynamics in our model is described by:

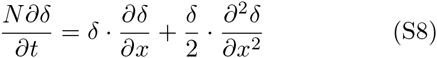

Compared to the traditional Burgers equation the diffusion coefficient is not constant but proportional to *δ*.

In KtW model the population collapses do not always happen in the order dictated by their relative sizes. When a collapse of a smaller population happens it only causes a small rescaling of populations, and the subsequent collapse of the largest population leads to a situation where these two populations are nearly equal in size. This dramatically increase the likelihood for further re-orderings between these two species, resulting in an intermittent dynamical period of fights for “dominance” between these two species.

The most likely “mistake” changing the order of collapses is when the second largest population “jumps the gun”and collapses ahead of the largest one. Repeating the above derivation in this case one gets *P*_2_(2)/*P*_1_(2) = 1 − *P*_2_(0) + *γ* ≃ 1 − *P*_2_(0). In the asymptotic case where populations are equidistantly distributed on the logarithmic scale one has *P_k_*(∞) = *γ*^(*k*−1)/*N*^ · (1 − *γ*^1/*N*^)/(1 − *γ*). In the limit where *γ*^1/*N*^ = *δ*\* ≪ 1 the top two populations occupy most of the carrying capacity and are approximately equal to 1 − *δ*\* and *δ*\* (up to the second order in *δ*\*). Hence, when the second largest population collapses ahead of the first one it leads to an instant and dramatic increase in *δ*_1_ to 1 − *P*_2_(0) = 1 − *δ*\* up from its steady state value of *δ*\*. The Eq. S7 describes the subsequent relaxation of *δ*_1_(*t*) back to *δ*\*. Indeed, if one disregards the first term in the r.h.s. of the equation but *δ*_1_ ≃ 1 one gets 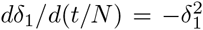. Hence initially the gaps starts relaxing as

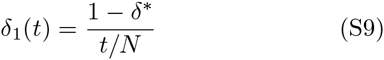

It takes about (1 − *δ*\*)/*δ*\* collapses for *δ*_1_(*t*) to get down close to *δ*\*. At this point one cannot completely disregard *δ*_2_ but one can still assume that it stays close to its the steady state value *δ*\* = *γ*^1/*N*^. In this case 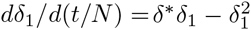 or

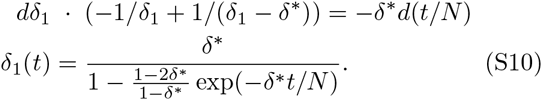

## Footnotes

[1] The ratio *δ_N_* for the currently smallest population *P_N_* is defined by its value after the next collapse when it becomes the second smallest.

